# Machine learning model interpretations explain T cell receptor binding

**DOI:** 10.1101/2023.08.15.553228

**Authors:** Brandon Carter, Jonathan Krog, Michael E. Birnbaum, David K. Gifford

**Author notes:** Correspondence to: David Gifford < >. Equal contribution (Brandon Carter, Jonathan Krog).

## Abstract

T cells mediate immune responses against pathogens and cancer through T cell receptors (TCRs) that recognize foreign peptides presented on the cell surface by Major Histocompatibility Complex (MHC) proteins. TCRs carry enormous diversity and differ across individuals, and mechanisms that determine TCR-pMHC binding are poorly understood. The ability to predict TCR-pMHC interactions would accelerate development of cellular therapeutics to design TCRs that specifically bind to a target of interest. We designed a randomized library of 19^6^ TCR CDR3*β* sequences and experimentally evaluated their affinities for the Tax-A02 peptide-MHC target. We trained ML models that predict TCR binding to Tax-A02 from TCR sequence and used model interpretation to identify TCR sequence features associated with binding. We found these features accurately mirror the true sequence features in our experimental data.

## 1. Introduction

T cells play an important role in the immune system’s defenses against pathogenic infection and cancer. T cells receptors (TCRs) recognize foreign peptide antigens presented on the cell surface by Major Histocompatibility Complex (MHC) proteins. TCRs exhibit moderate affinity for their targets, and a given TCR may recognize multiple MHC-displayed antigens, permitting recognition of a nearly unlimited complement of antigens that may arise from pathogenic infection or cancer (Mason, 1998). Each individual has a diverse repertoire of T cells whose TCR specificities for cognate pMHCs are determined by random V(D)J recombination during lymphocyte maturation.

Each TCR is a heterodimer comprising an *α* and *β* chain. Each chain consists of three loops, called CDR1, 2, and 3. The CDR1 and 2 loops contact the MHC helices, while the CDR3 loops primarily interact with the peptide (Garcia & Adams, 2005). CDR3 loops carry significant sequence diversity to permit TCR binding to a wide array of cognate antigens. While it has been possible to identify the range of antigens recognized by a single TCR (Birnbaum et al., 2014), or to identify which TCRs in a T cell repertoire recognize a specific antigen of interest, it has been more difficult to screen many TCRs against many potential targets.

Accurate prediction of TCR-pMHC binding would allow better prediction of responses to immunotherapies and personalized design of more effective T cell vaccines. These models can also be applied to design novel TCRs that bind antigens of interest, for instance in cellular therapies for cancer. However, this problem is challenging as a result of the large input space of possible TCR sequences × MHC allelic variants × target antigens that is presently sparsely sampled. Bulk CDR3 sequence data available in public repositories is limited and has been demonstrated to be generally poor quality that has hindered machine learning (ML) model accuracy (Montemurro et al., 2021). While prior efforts have studied general models that predict binding between any TCR sequence and multiple targets using the available paired TCR-pMHC data (Montemurro et al., 2021; Jokinen et al., 2023; Lu et al., 2021), here we focus on training models on densely sampled TCR CDR3*β* sequence mutations to predict binding to specific targets and reveal underlying binding mechanisms through model interpretation.

We trained ML models to predict binding affinity between mutated A6 TCR CDR3*β* sequences and the Tax peptide (LLFGYPVYV) presented by HLA-A*02:01, a complex recognized by the well-studied A6 TCR (Garboczi et al., 1996; Ding et al., 1999). We synthesized a library of TCRs that randomly mutated six amino acids of the A6 CDR3*β* chain that lie within the TCR-pMHC binding interface. The resulting modified TCRs were expressed in a yeast system and screened against the Tax-A02 complex to experimentally measure binding enrichment after three selection rounds. After each selection round the selected TCRs were sequenced.

Our experimental data offers the benefit of dense sampling of mutations to the peptide-binding region of the CDR3*β* loop to measure how each mutation maintains, eliminates, or enhances binding to the target. We trained ML models on these data and found neural networks outperform baseline methods on held-out experimental data. We interpreted our models, and the explanations reveled that the models classify TCR sequences based upon biologically meaningful sequence features present in the experimental data.

Our contributions include:

1. We designed a randomized, dense library of mutant A6 TCR sequences and experimentally measured the affinities of mutant TCRs against the Tax-A02 target.
2. We train ML models that can predict whether a TCR mutant sequence binds Tax-A02.
3. We show how model interpretation can be used to (1) evaluate whether the model has learned true biological mechanisms and (2) expose important TCR-pMHC binding residues that can be evaluated in future studies.

## 2. Methods

### 2.1 Experimental protocol and randomized TCR library design

The data described in this work was generated with consecutive rounds of high-throughput binding assays (“selections”) which utilize diversified libraries of T cell receptors (TCRs) selected against a peptide-MHC (pMHC) target of interest (Figure 1). The TCR framework used herein is a patient derived clone that has been previously modified (described in Aggen et al. (2011)) for stability when displayed on the surface of yeast cells.

**Figure 1.**
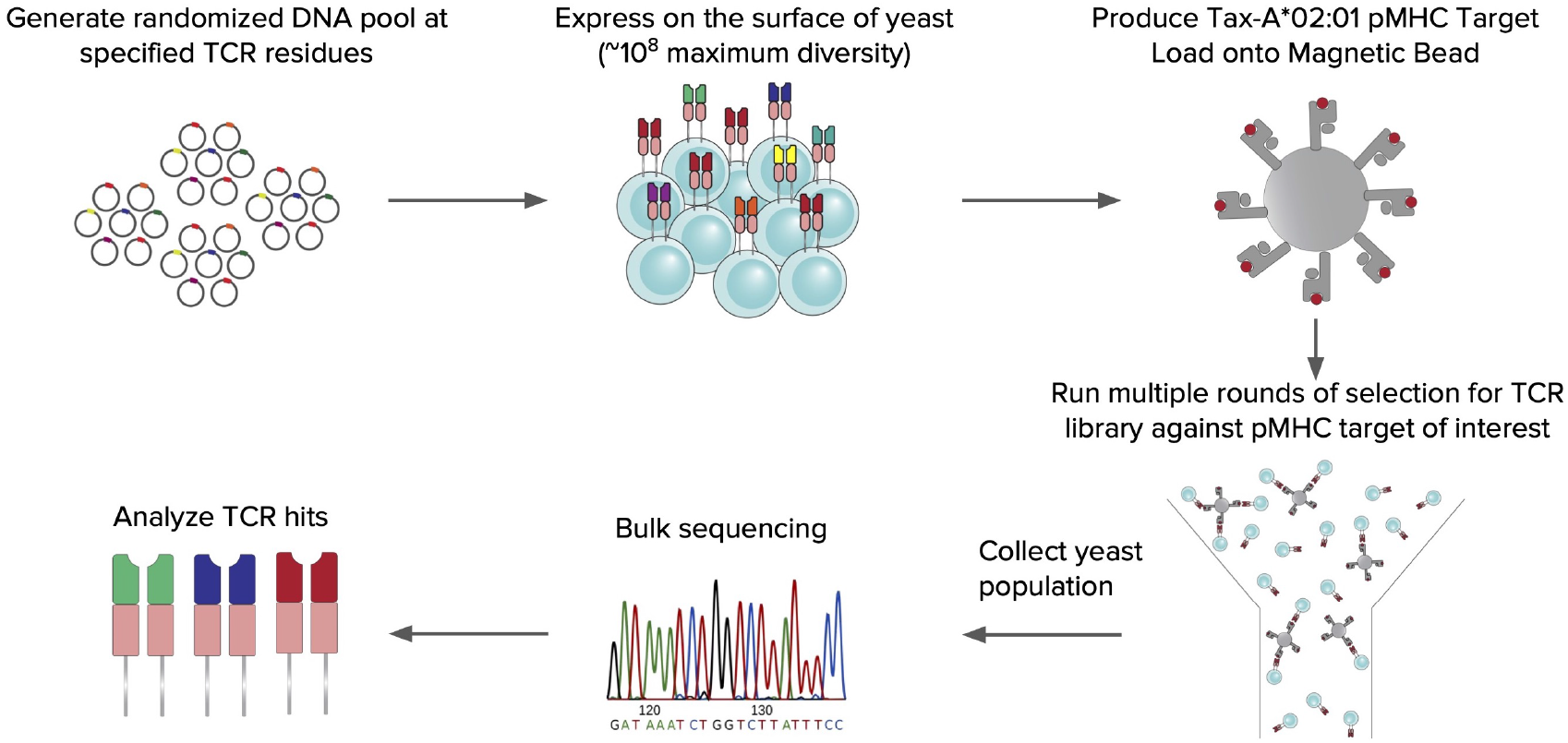
Overview of experimental protocol for generating TCR-pMHC binding data with yeast display selections. The workflow includes producing DNA variants for TCRs of interest, expressing these TCR constructs on the surface of yeast, using magnetic beads with pMHC loaded on the surface to separate binders and then identifying corresponding TCRs via deep sequencing.

**Figure 2.**
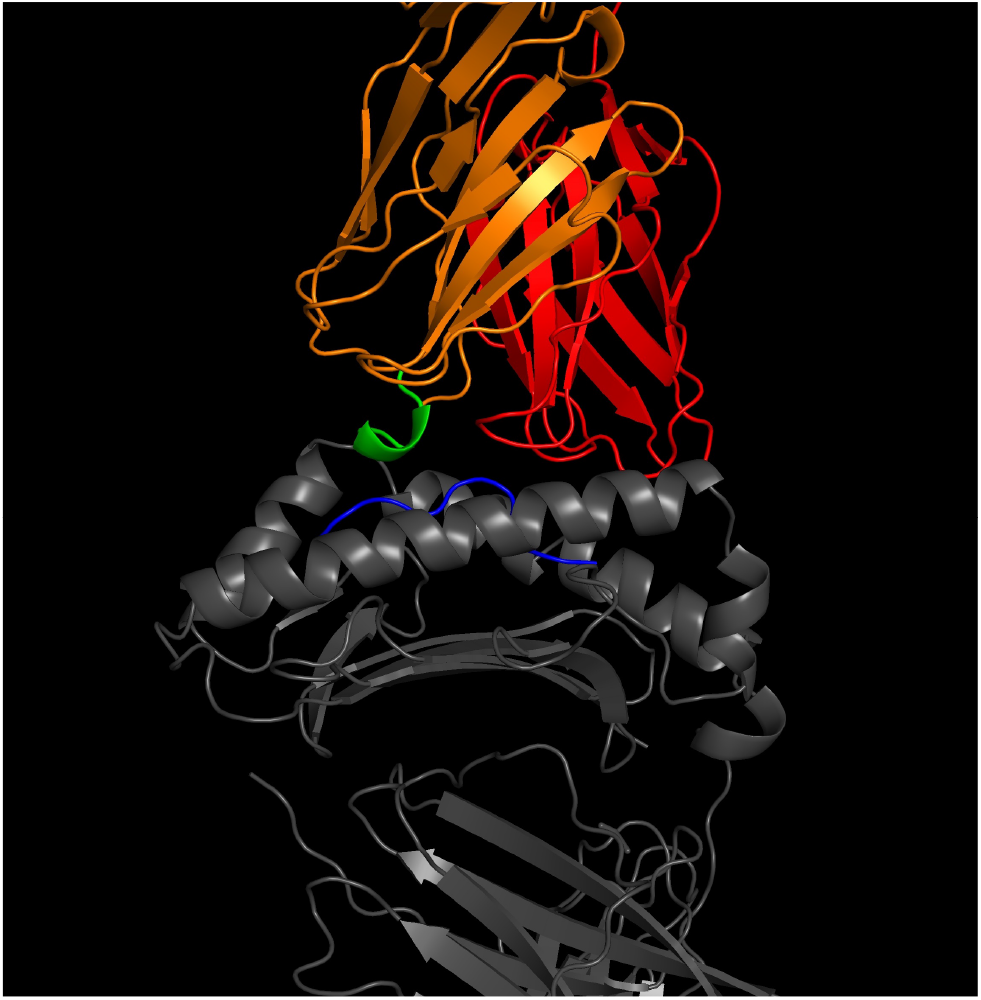
Crystal structure of A6 TCR (CDR3*α* red, CDR3*β* orange), Tax peptide (blue), and HLA-A*02:01 (gray) complex from Garboczi et al. (1996) (PDB: 1AO7). Our TCR library design mutated six amino acid residues in the CDR3*β* chain that are closest to the peptide of the bound pMHC structure (green, see Section 2.1). Visualization was created using PyMOL (Schrödinger, 2021).

Using this stability enhanced framework as a starting point, we then generated our TCR library by selecting six amino acid residues in the CDR3*β* chain that make closest contact with the peptide of the bound pMHC structure. The diversity at these positions was introduced using randomized primers from Ella Biotech that have a defined and uniform distribution of amino acids at each position, without cysteine. Once created, these randomized DNA inserts along with the pCT302 yeast display vector were electroporated into the yeast strain RJY100 and assembled via homologous recombination resulting in a theoretical diversity of 19^6^ unique TCR variants.

This TCR library was then used in three consecutive rounds of magnetic-activated cell sorting (MACS) selections against the pMHC complex of Tax–HLA-A*02:01 to enrich for TCR variants that confer enhanced binding to the target complex. Resulting yeast populations after each round of selection were deeply sequenced and the generated data is utilized in the following ML applications.

Our dataset includes the read count of each unique CDR3*β* sequence in the randomized library (R0) as well as after the three selection rounds (R1–R3). We removed sequence reads whose mutated CDR3*β* sequence was not 6 amino acids due to sequencing errors. The final dataset contains 208784 unique TCR CDR3*β* sequences (details in Appendix C).

We randomly split our dataset into 90% train/val and 10% test sets. The train/val set was further split using 10-fold cross-validation for model tuning. After validation, the final models were trained on the entire train/val set before evaluation on the held-out test set.

We define a classification task to predict whether a TCR sequence is *enriched* (≥10 reads observed in R3) for binding to Tax-A02 (additional experiments in Appendix C). The full dataset contains 731 enriched sequences (654 in train/val set, 77 in test set). The wild-type A6 TCR (`PGLAGG`) had 24 reads observed in R3.

### 2.2 Machine learning models

We train neural networks to predict TCR sequence enrichment for binding Tax-A02 from its mutated CDR3*β* sequence. The six mutated amino acids are embedded using BLOSUM50 vectors (Henikoff & Henikoff, 1992). Each neural network contains two hidden layers with 500 neurons and ReLU activation and outputs the probability of enrichment. We minimize cross-entropy loss using the Adam optimizer (Kingma & Ba, 2015). Additional implementation details are described in Appendix A.

To address the significant class imbalance in our dataset (0.35% positives), we randomly upsampled positive examples in the training set (1:10 positive:negative ratio). We generally found that replicate neural networks with different random initialization exhibited significant variation in performance (Figure A.1). As a result, our final TCREnsemble model is an ensemble of 10 replicate neural networks trained with different random initialization to mitigate idiosyncratic errors by individual networks. This ensemble outputs the mean prediction of the member networks. We found that our ensemble generally improved performance over individual networks through cross-validation (Figure A.1).

We trained *k*-nearest neighbors (*k*-NN) and logistic regression classifiers as baseline control models. These models take as input the same BLOSUM50 embeddings as the neural networks. We found *k* = 8 nearest neighbors to maximize cross-validation performance (details in Appendix A).

### Model interpretation

To identify the sequence features learned by the ML models, we adopt the Sufficient Input Subsets (SIS) method for black-box model interpretation (Carter et al., 2019; 2020). For a given sequence predicted to be enriched by our model, SIS identifies a minimal subset of amino acids in the sequence whose values alone suffice for the model to predict enrichment with high confidence (here ≥ 90% probability). The non-SIS amino acids are masked using the `X` (unknown) amino acid as represented by BLOSUM50. The SIS can be understood as the support for the model’s decision on a particular instance.

We can also cluster SIS explanations for many sequences predicted to the enriched by the model to identify global feature patterns the model has learned to associate with sequence enrichment. Here, SIS are clustered using DB-SCAN (Ester et al., 1996) with Levenshtein (edit) distance between masked sequences.

## 3. Results

### 3.1 TCREnsemble predicts enriched TCR sequences

Table 1 shows the evaluation of ML models that predict enrichment of TCR CDR3*β* sequences for the Tax-A02 target. Our neural network TCREnsemble model outperformed logistic regression and *k*-nearest neighbors baselines on held-out test sequences from our experimental data.

**Table 1.** Performance of ML models at classifying enriched TCR sequences in held-out test set. Performance of TCREnsemble and individual ensemble members is given as mean ± standard deviation over five model replicates with different random initialization. The last column shows the number of predicted positive sequences in the test data (test set contains 20878 sequences with 77 true positives).

|  | Accuracy | AUROC | Avg. Precision | Precision | Recall | F1 | Pred. Positive |
| --- | --- | --- | --- | --- | --- | --- | --- |
| TCREnsemble | 99.7% ± 0.0% | 0.985 ± 0.0 | 57.3% ± 0.7% | 52.7% ± 0.5% | 82.3% ± 2.2% | 64.2% ± 0.8% | 120.4 ± 3.4 |
| Individual Ensemble Members | 99.7% ± 0.0% | 0.985 ± 0.0 | 57.5% ± 1.8% | 52.0% ± 2.5% | 77.4% ± 5.0% | 62.1% ± 2.7% | 114.8 ± 8.7 |
| Logistic Regression | 99.6% | 0.974 | 37.8% | 44.4% | 20.8% | 28.3% | 36 |
| *k*-Nearest Neighbors | 99.6% | 0.965 | 43.9% | 49.3% | 42.9% | 45.8% | 67 |

To interpret the sequences features used by TCREnsemble to classify TCR sequences, we applied SIS (Section 2.3) to find minimal subsets of TCR sequences whose values alone permit the model to predict TCR enrichment with at least 90% confidence. Figure 3 shows the result of clustering the SIS explanations found to explain test set TCR sequences predicted to be enriched for Tax-A02 (all explanations shown in Table B.1). These clusters indicate patterns of amino acids that the model has learned to associate with binding Tax-A02, even without information about the masked residues, and thus may provide hypotheses about residues important for TCR binding to the Tax-A02 complex.

**Figure 3.**
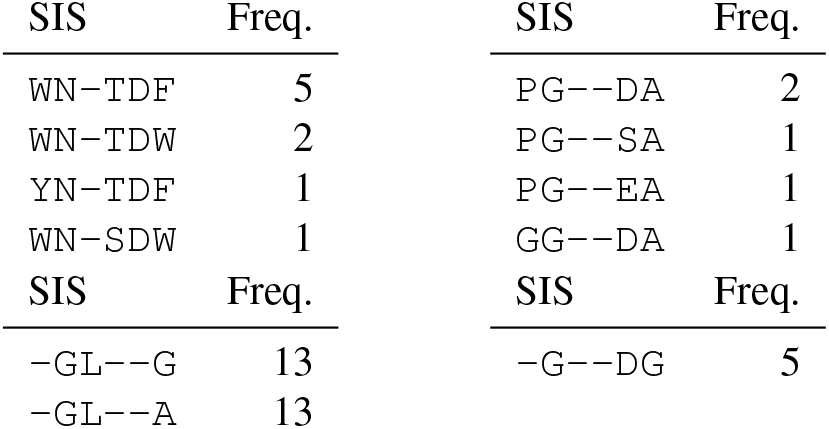
Four clusters of sufficient input subset (SIS) explanations found across TCR sequences predicted to be enriched for Tax-A02 by our TCREnsemble model. Each cluster contains a SIS explanation and frequency indicating the number of TCR sequences in the test data whose prediction was explained with that SIS. All SIS shown here are sufficient for confident (≥ 90%) enrichment prediction by the ML model. Individual SIS are shown in Table B.1.

### 3.2 Model interpretations mirror biological sequence features

To evaluate whether our ML model operates based upon biologically meaningful sequence features, we compared the SIS explanations from the model to amino acids that carry high information content for enriched sequences from our ground-truth experimental data. Figure 4 shows sequence logos for (A) ground-truth enriched TCR sequences in our full dataset (731 sequences) and (B) SIS explanations for test sequences predicted to be enriched by the ML model (81 SIS). We find that high information content residues in SIS explanations mirror high information content residues in ground-truth enriched sequences (*p* = 0.001 for Jensen-Shannon divergence between the ground-truth enriched and SIS distributions vs. bootstrapped test sequences, see Appendix B). Thus, we find significant evidence that our ML model is classifying TCR sequences using biologically meaningful features.

**Figure 4.**
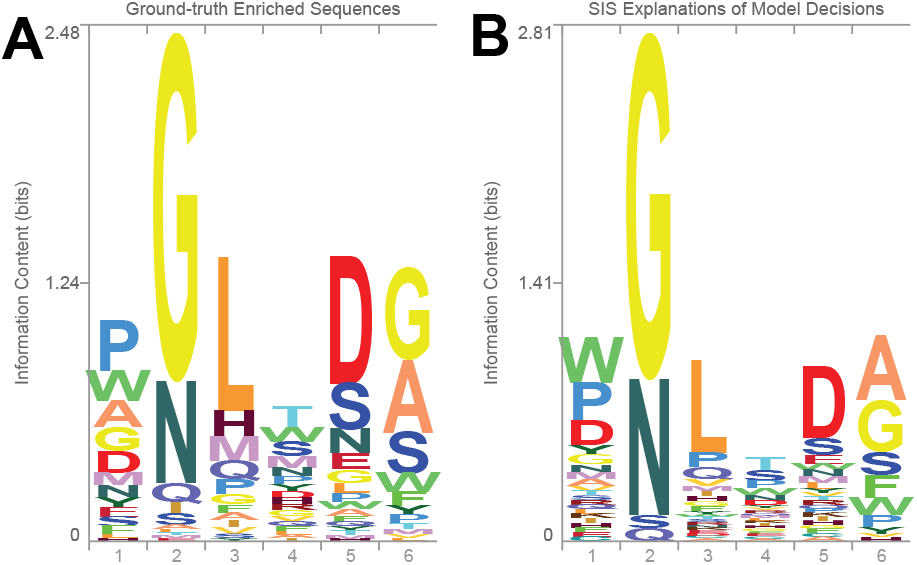
Sequence logos for (A) ground-truth enriched TCR sequences in the full dataset and (B) sufficient input subset (SIS) explanations of decisions by our TCREnsemble model for predicted enriched sequences in the test set.

## 4. Discussion

We showed that the interpretation of ML models trained on experimental TCR-pMHC binding data reveals mechanisms of T cell receptor binding. Our models are trained on experimental data we collected from a library of randomized mutations to the CDR3*β* loop of the A6 TCR. We found that sequence features learned by our ML model matched sequence features with high information content in the experimental data. One challenge of training models on TCR-pMHC binding data is the significant class imbalance, and we found upsampling enriched sequences was sufficient to train neural network models. Another challenge faced by previous work training TCR-pMHC prediction models is the lack of negative TCR-pMHC pairs (Montemurro et al., 2021). By screening a large randomized TCR library, our dataset includes experimentally validated negative binders.

One application of our ML models is optimization of TCR sequences with superior binding affinity to previously observed sequences, for instance to design therapeutic T cells that target tumor cells. In addition, our methods can be extended to predict both the *affinity* and *specificity* of TCR sequences to multiple targets to design TCRs that are target-specific and avoid off-target interactions (Saksena et al., 2022). We are exploring both directions in ongoing work.

## A. Details of Machine Learning Models

### Model Architecture Details

Our machine learning models take as input embedded sequences specifying the amino acid identities of the six mutated TCR CDR3*β* sequence positions. Our final models use BLOSUM50 encodings, and we also considered one-hot encodings that were not found to be significantly superior for our TCREnsemble model (results shown in Table A.1). Given each amino acid sequence is represented by a 20-dimensional BLOSUM50 vector, the full input is a flattened 120-dimensional vector.

Our neural network architecture contains two hidden layers each with 500 dimensions and ReLU activation and outputs the probability of enrichment (softmax). All models are trained for 10 epochs with a batch size of 128. We minimize cross-entropy with the Adam optimizer (Kingma & Ba, 2015) with a learning rate of 0.0001 and weight decay of 0.0005. Positive sequences in the training set are randomly upsampled 1:10 positive:negative to decrease the effect of imbalanced data.

We used 10-fold cross-validation of the train/val set to evaluate the following hyperparameters: learning rate ∈ [0.0001, 0.001, 0.01], hidden dimension ∈ [50, 100, 500], batch size ∈ [128, 256, 512], weight decay ∈ [0.0005, 0.005], encoding ∈ [BLOSUM50, one-hot], and positive:negative upsampling ratio ∈ [0.1, 0.25, 0.5, 1.0]. In general, we observed significant variation in model performance resulting from (1) random initialization of models on a given train/val fold, and (2) variation across different train/val folds, and we found multiple configurations of the model hyperparameters that worked well on our data. Figure A.1 shows the performance of 10 replicate neural networks (black points) trained with the same hyperparameters but different random initializations. To mitigate the errors made by individual models, we introduced an ensemble (TCREnsemble, red points in Figure A.1) that outputs the average prediction of over the 10 replicate networks (the arithmetic mean of their logits), and we found that the ensemble generally improved model performance over individual models.

**Figure A.1.**
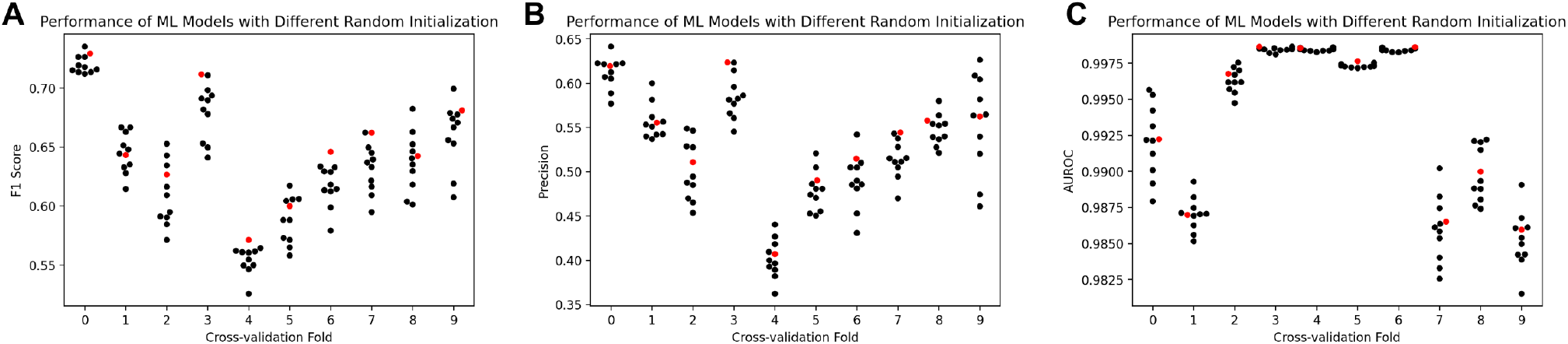
Performance of 10 replicate neural networks (black points) trained with the same hyperparameters but different random initialization evaluated by (A) F1 score, (B) Precision, and (C) AUROC on 10-fold cross-validation. The performance of an ensemble that outputs the mean prediction of the 10 individual networks is shown in red.

**Table A.1.**
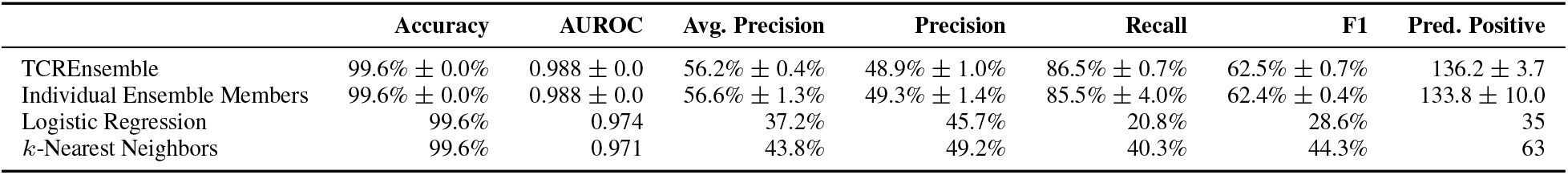
Test set performance of machine learning models with input TCR sequences encoded as one-hot vectors. Performance of TCREnsemble and individual ensemble members is given as mean ± standard deviation over five model replicates with different random initialization. The last column shows the number of predicted positive sequences in the test data (test set contains 20878 sequences with 77 true positives). Model performance with BLOSUM50 encodings of TCR sequences is shown in Table 1. Models were not further tuned and use the same hyperparameters as the models in Table 1 with BLOSUM50 encodings.

### Baseline Models

For our *k*-nearest neighbors baseline, we tuned hyperparameters including *k ∈* [1, …, 20], weight function [uniform, distance] (where distance weights points by the inverse of their distance), and positive:negative upsampling ratio [0, 0.1]. Neighbors were computed using Euclidean distance between BLOSUM50 encodings of TCR sequences. We found *k* = 8, distance weighting, and no upsampling of positives in the training set maximized F1 score on cross-validation data.

For our logistic regression baseline with *L*_2_ regularization, we tuned the regularization strength *C* ∈ [0.001, 0.01, 0.1, 1, 10, 100, 1000] (where *C* is the inverse of the regularization strength) and positive:negative upsampling ratio [0, 0.1]. We found *C* = 100 and no upsampling of positives maximized F1 score on cross-validation data.

Unlike training neural networks, for our baseline models, we found that not upsampling positive examples in the training set maximized model performance.

Our neural network models were implemented in PyTorch (Paszke et al., 2019), and our *k*-nearest neighbors and logistic regression models were implemented using scikit-learn (Pedregosa et al., 2011).

## B. Details of Model Interpretation

### Additional SIS Examples

Table B.1 shows the sufficient input subset (SIS) explanations found for all 81 test set sequences predicted by the ML model to be enriched with at least 90% confidence (SIS threshold 0.9). We mask sequences using the BLOSUM50 vector for the unknown amino acid (`X`) and note that the predicted confidence for enrichment of the fully masked sequence (`XXXXXX`) is 0.0013 (not enriched).

Sequence logos were created using Skylign (Wheeler et al., 2014).

### Comparing Distributions of Sequences

To measure the divergence between two distributions of sequences of length *n*, we first represent each distribution as a position probability matrix (PPM) that specifies the probability of each possible amino acid *i* occurring at each sequence position *j* assuming independence between positions, estimated using the observed sequences. For two sequence distributions *P* and *Q*, we then compute the Jensen-Shannon divergence at each sequence position *j* ∈ {1, …, *n*}:

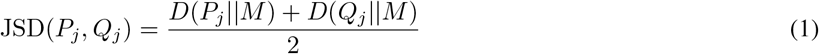

where *D* is the Kullback-Leibler divergence and 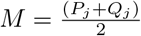. Then, we define the final divergence between *P* and *Q* as the sum of the Jensen-Shannon divergence over the sequence positions:

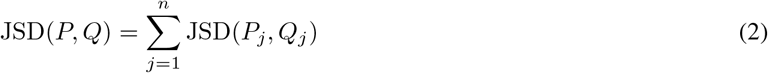

Here, we measure the divergence between two distributions of sequences: (1) TCR sequences defined as enriched in our ground-truth experimental dataset and (2) SIS explanations of test set TCR sequences predicted enriched by our ML model. To map SIS explanations into a PPM, we replace masked (non-SIS) sequence positions with the `X` (unknown) amino acid, and define `X` as a uniform distribution over the 20 amino acids.

To compute whether the divergence between the two sequence distributions above is statistically significant, we estimate the distribution of divergence between the ground-truth enriched sequences and randomly drawn test set sequences (of any class) using bootstrapping and the Monte Carlo test (Davison & Hinkley, 1997). We compute *R* = 1000 bootstrapped samples of test sequences (each sample containing the same number of sequences as we have SIS explanations) and estimate the one-sided Monte Carlo *p*-value as:

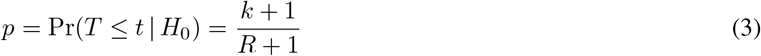

where *t* is the observed divergence between the ground-truth and SIS sequence distributions, *T* is the distribution of divergence between the ground-truth enriched sequences and randomly drawn sequences under the null hypothesis *H*_0_, and *k* counts the number of simulated values *t*^∗^ ∼ *T* where *t*^∗^ ≤*t*. Figure B.1 shows an example sequence logo for 81 sequences from the test set drawn at random. Empirically, we found the divergence between ground-truth and SIS sequences to be 0.24 and the divergence between ground-truth and random test sequences to be 1.39 *±* 0.08.

**Figure B.1.**
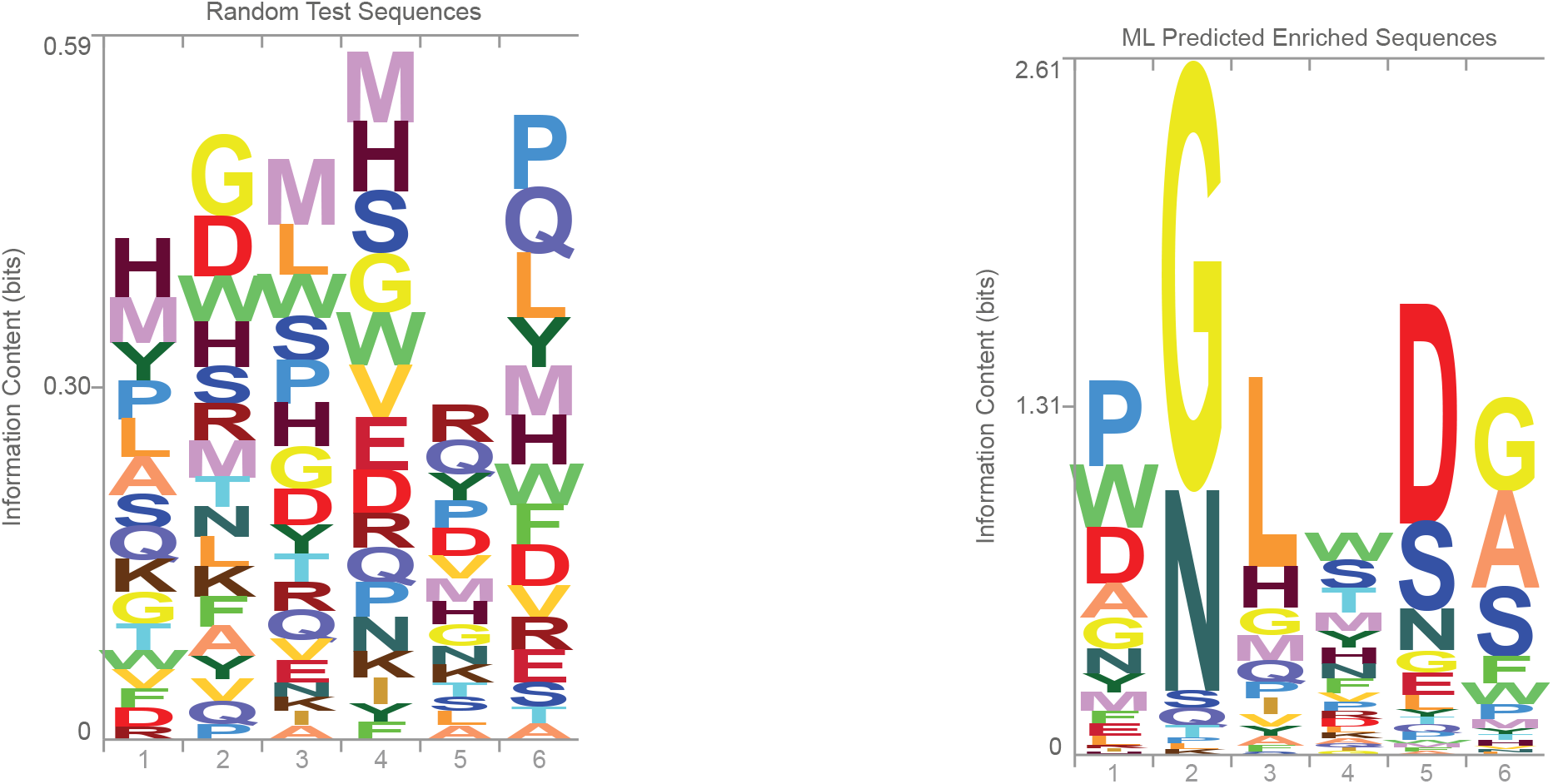
Sequence logo for 81 sequences randomly selected from the test set (note difference in y-axis scale compared to Figure 4).

**Figure B.2.**
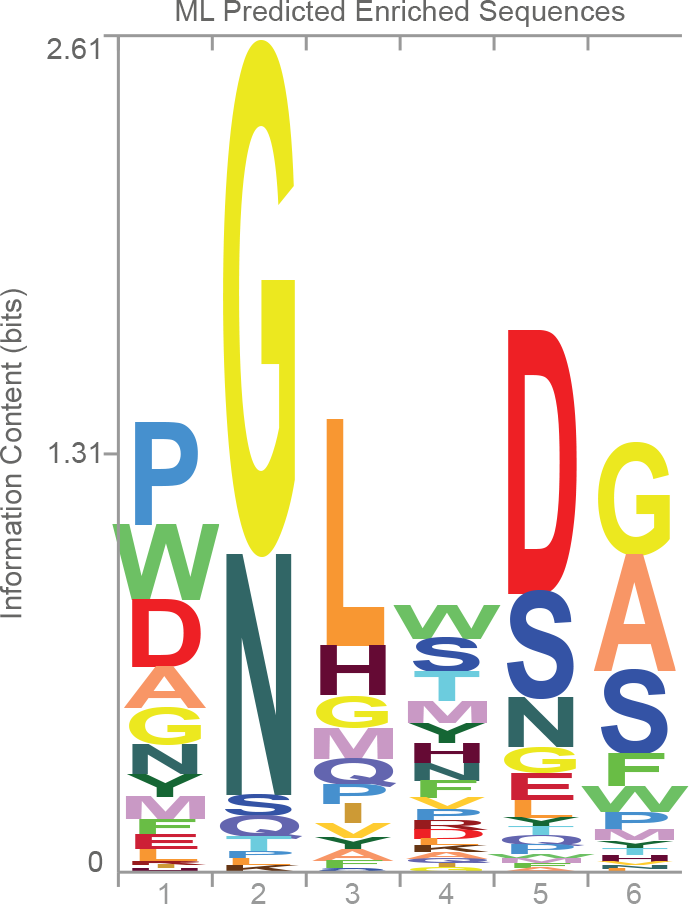
Sequence logo for ML predicted enriched sequences in the test set.

Figure B.2 shows the sequence logo for all 120 test sequences predicted to be enriched by the ML model. We found the divergence between the distribution of predicted positive test sequences and ground-truth sequences was 0.12 and significantly lower than divergence between the ground-truth and random test sequences (*p* = 0.001 vs. randomly drawn test sequences).

**Table B.1.**
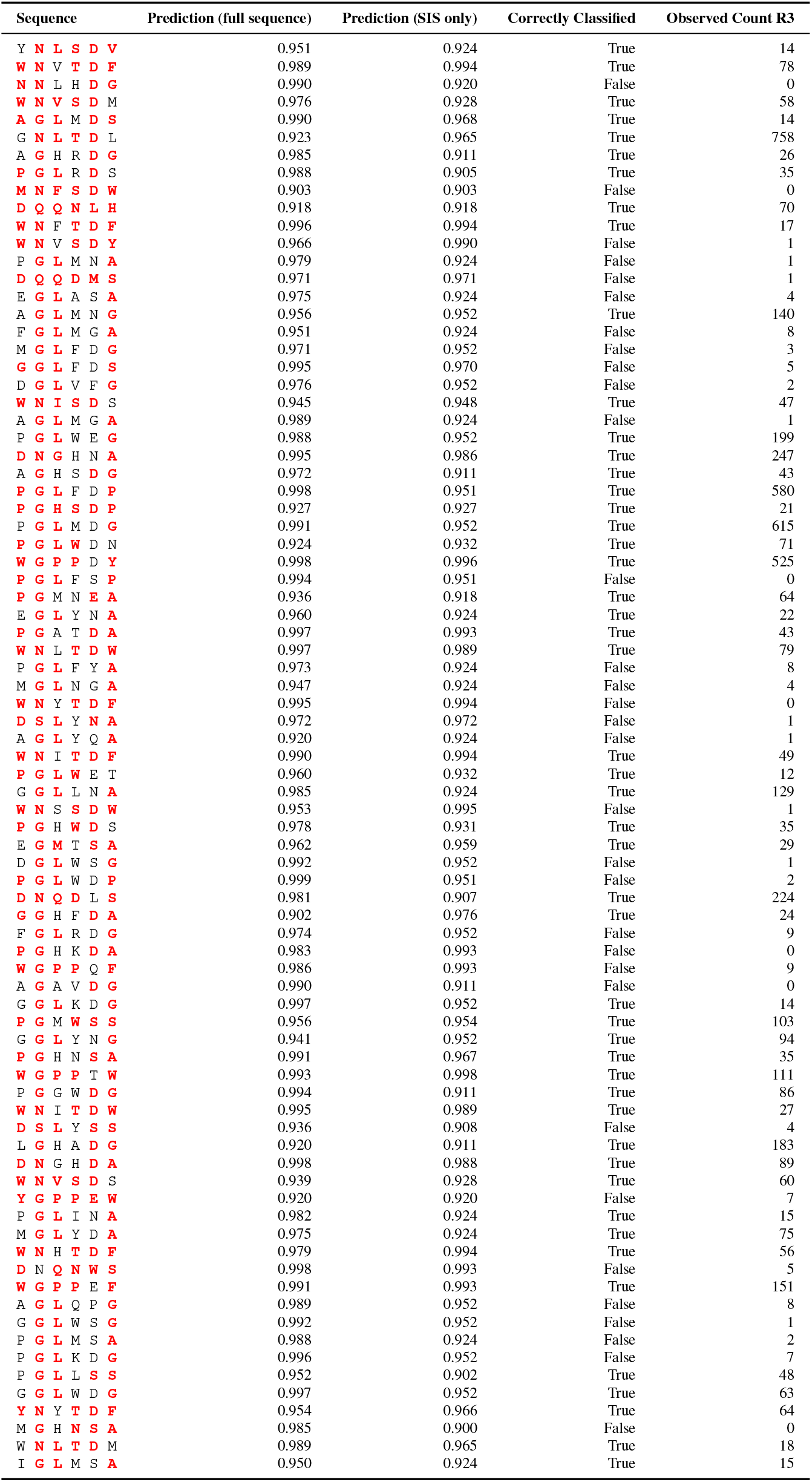
Sufficient input subset (SIS) explanations found to interpret decisions of all test set TCR sequences predicted to be enriched with at least 90% confidence by the ML model (SIS threshold 0.9). SIS residues are highlighted in red. Predictions give the model confidence of TCR enrichment on either the full sequence or SIS residues only. For prediction on SIS only, non-SIS residues are replaced with X (Section 2.3). The fourth column specifies if the full sequence is correctly classified by the model.

## C. Dataset Details and Sensitivity to Enrichment Threshold

As described in Section 2.1, we used a threshold of 10 reads in R3 to define enriched (positive) TCR sequences in our dataset (results in Section 3). Table C.1 shows the count of unique TCR sequences and read count distributions observed for each selection round (Section 2.1). Figure C.1 shows the number of enriched sequences in the full dataset as a function of the R3 read count threshold used to define enrichment.

**Table C.1.**
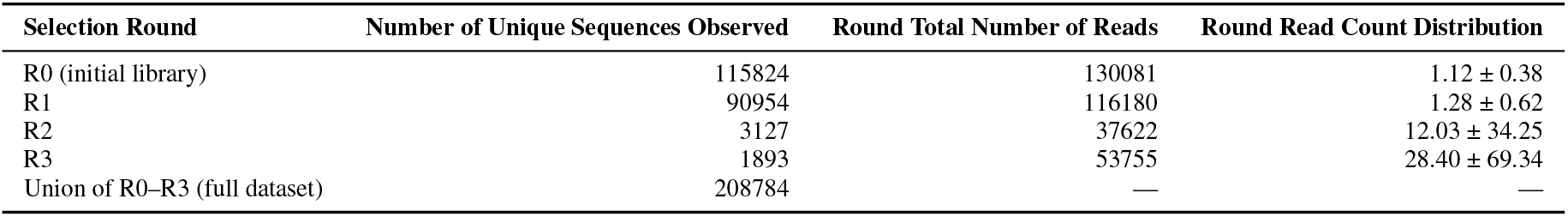
Number of unique TCR sequences and read counts observed after sequencing the results of each selection round and removing sequencing errors (Section 2.1). Round read count distributions specify the mean ±standard deviation of read counts for each unique sequence observed in each round.

| Selection Round | Number of Unique Sequences Observed | Round Total Number of Reads | Round Read Count Distribution |
| --- | --- | --- | --- |
| R0 (initial library) | 115824 | 130081 | $1.12 \pm 0.38$ |
| R1 | 90954 | 116180 | $1.28 \pm 0.62$ |
| R2 | 3127 | 37622 | $12.03 \pm 34.25$ |
| R3 | 1893 | 53755 | $28.40 \pm 69.34$ |
| Union of R0–R3 (full dataset) | 208784 | — | — |

**Figure C.1.**
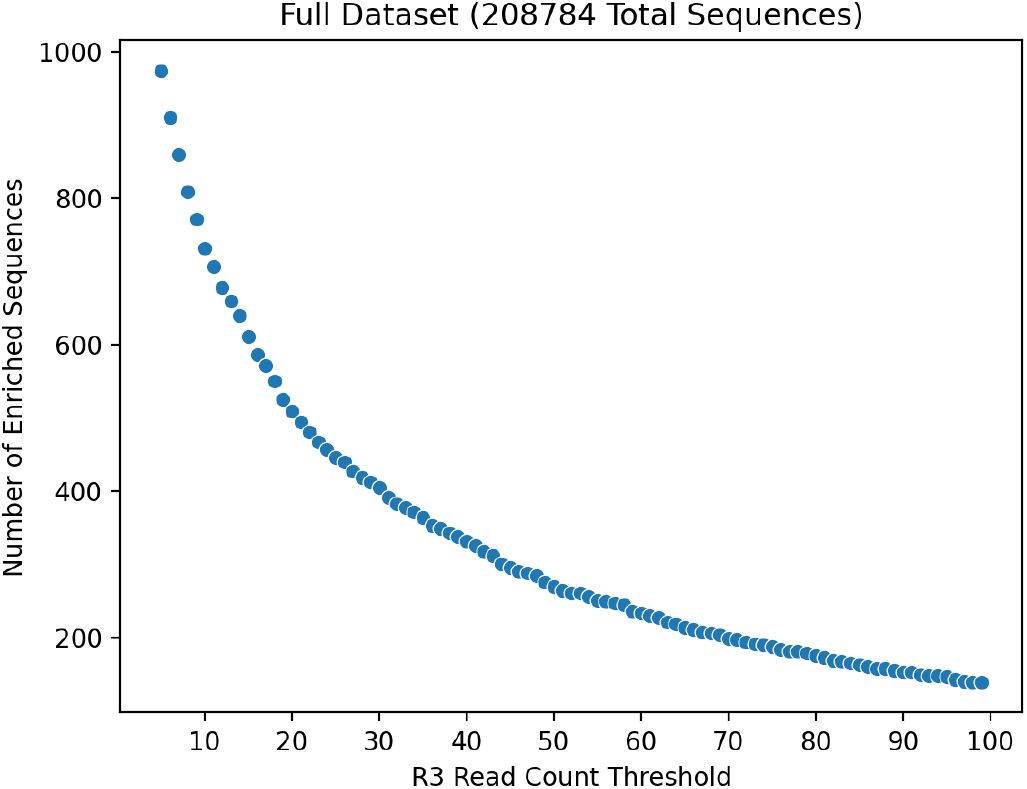
Number of enriched sequences in the full dataset at different thresholds for R3 read count.

Table C.2 reports model performance results of training and evaluating our models where positive sequences are defined to have ≥20 reads in R3 (454 enriched sequences in training set, 55 enriched sequences in test set). These models were trained on the full train/val set and tested on the held-out test set (same sequences as Table 1 with a stricter threshold for defining positives). These models were trained using the same parameters as the models in Table 1 and were not separately tuned. We found our TCREnsemble model outperformed the baseline methods.

**Table C.2.**
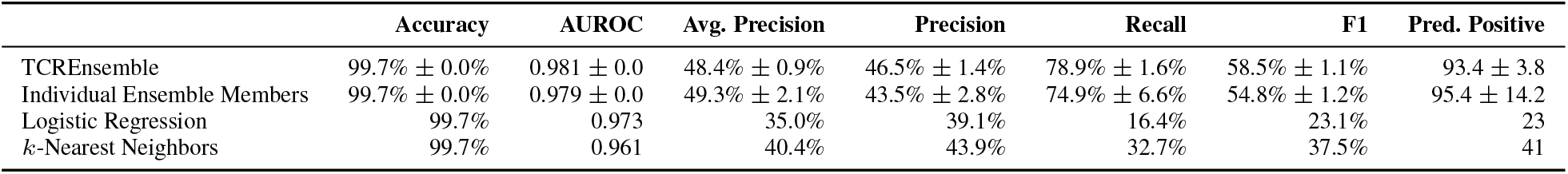
Test set performance of machine learning models using enrichment threshold of 20 reads in R3. Performance of TCREnsemble and individual ensemble members is given as mean ±standard deviation over five model replicates with different random initialization. The last column shows the number of predicted positive sequences in the test data (test set binarized using R3 read count of 20 contains 20878 sequences with 55 true positives). Models were not further tuned and use the same hyperparameters as the models in Table 1.

